# A microbially derived tyrosine sulfated peptide mimics a plant peptide hormone

**DOI:** 10.1101/116897

**Authors:** Rory N. Pruitt, Anna Joe, Weiguo Zhang, Wei Feng, Valley Stewart, Benjamin Schwessinger, José R. Dinneny, Pamela C. Ronald

## Abstract

- The biotrophic pathogen *Xanthomonas oryzae* pv. *oryzae* (*Xoo*) produces a sulfated peptide named RaxX, which shares similarity to peptides in the PSY (*p*lant peptide containing *s*ulfated *t*yrosine) family. We hypothesize that RaxX functionally mimics the growth stimulating activity of PSY peptides.
- Root length was measured in Arabidopsis and rice treated with synthetic RaxX peptides. We also used comparative genomic analysis and Reactive Oxygen Species (ROS) burst assay to evaluate the activity of RaxX and PSY peptides.
- Here we found that a synthetic sulfated RaxX derivative comprising 13 residues (RaxX13-sY), highly conserved between RaxX and PSY, induces root growth in Arabidopsis and rice in a manner similar to that triggered by PSY. We identified residues that are required for activation of immunity mediated by the rice XA21 receptor but that are not essential for root growth induced by PSY. Finally, we showed that a *Xanthomonas* strain lacking *raxX* is impaired in virulence.
- These findings suggest that RaxX serves as a molecular mimic of PSY peptides to facilitate *Xoo* infection and that XA21 has evolved the ability to recognize and respond specifically to the microbial form of the peptide.

## Introduction

Some plant and animal pathogens employ molecular mimicry to gain evolutionary advantages (Mitchum *et al.,* 2012). Such microbial molecules include those that mimic ligands of host receptors, substrates of host enzymes, or host proteins themselves (Knodler *et al.,* 2001; Nesic *et al.,* 2010). Some plant pathogens produce small molecules that mimic plant hormones required for growth, development and regulation of innate immunity.

A well-studied case of hormone mimicry in plants is the production of coronatine by the gram-negative biotrophic bacterium *Pseudomonas syringae* (Weiler *et al.,* 1994). Coronatine structurally and functionally mimics jasmonoyl-L-isoleucine (JA-Ile), a bioactive form of the plant hormone jasmonic acid (JA) (Weiler *et al.,* 1994). JA positively regulates defense against chewing insects and necrotrophic pathogens and negatively regulates defense against biotrophic and hemibiotrophic pathogens. Coronatine produced during *P. syringae* infection mimics JA action, suppressing the host defense response.

Plant parasitic nematodes and fungi also produce mimics of endogenous plant hormones. For example, nematodes produce peptides similar to plant CLAVATA3/ESR (CLE) peptides (Chen *et al.,* 2015), which regulate shoot meristem differentiation, root growth, and vascular development. Nematode CLEs are secreted into plant tissues where they induce specific host cells to differentiate into feeding cells that benefit the parasite (Wang *et al.,* 2005; Mitchum *et al.,* 2008; Yamaguchi *et al.,* 2016). Another example is C-TERMINALLY ENCODED PEPTIDEs (CEPs), a large and diverse family of effector peptides produced by sedentary plant-parasitic nematodes (PPNs). Plant CEPs inhibit root growth and increase the gene expression of a nitrogen transporter in response to nitrogen starvation. It is hypothesized that the parasite produced CEPs promote nitrogen uptake and reduce the size of the feeding site where the PPNs maintain biotrophic interactions (Eves-Van Den Akker *et al.,* 2016). Finally, the root-infecting fungus *Fusarium oxysporum* secretes a functional mimic of plant regulatory peptide RALF (*r*apid *al*kalinization *f*actor). RALF from *Fusarium oxysporum* induces extracellular alkalinization in the host apoplast which favors pathogen multiplication (Murphy & De Smet, 2014; Masachis *et al.,* 2016).

We have recently shown that the rice receptor XA21 is activated by a sulfated protein, called RaxX, produced by the bacterial pathogen *Xanthomonas oryzae* pv. *oryzae* (*Xoo*). RaxX triggers a robust and effective immune response in rice expressing XA21 (Song *et al.,* 1995;

Pruitt *et al.,* 2015). A synthetic 21-amino acid sulfated derivative of RaxX (RaxX21-sY) from *Xoo* strain PXO99 (Fig. 1A) is sufficient to activate XA21-mediated immune responses (Pruitt *et al.,* 2015).

**Figure 1.**
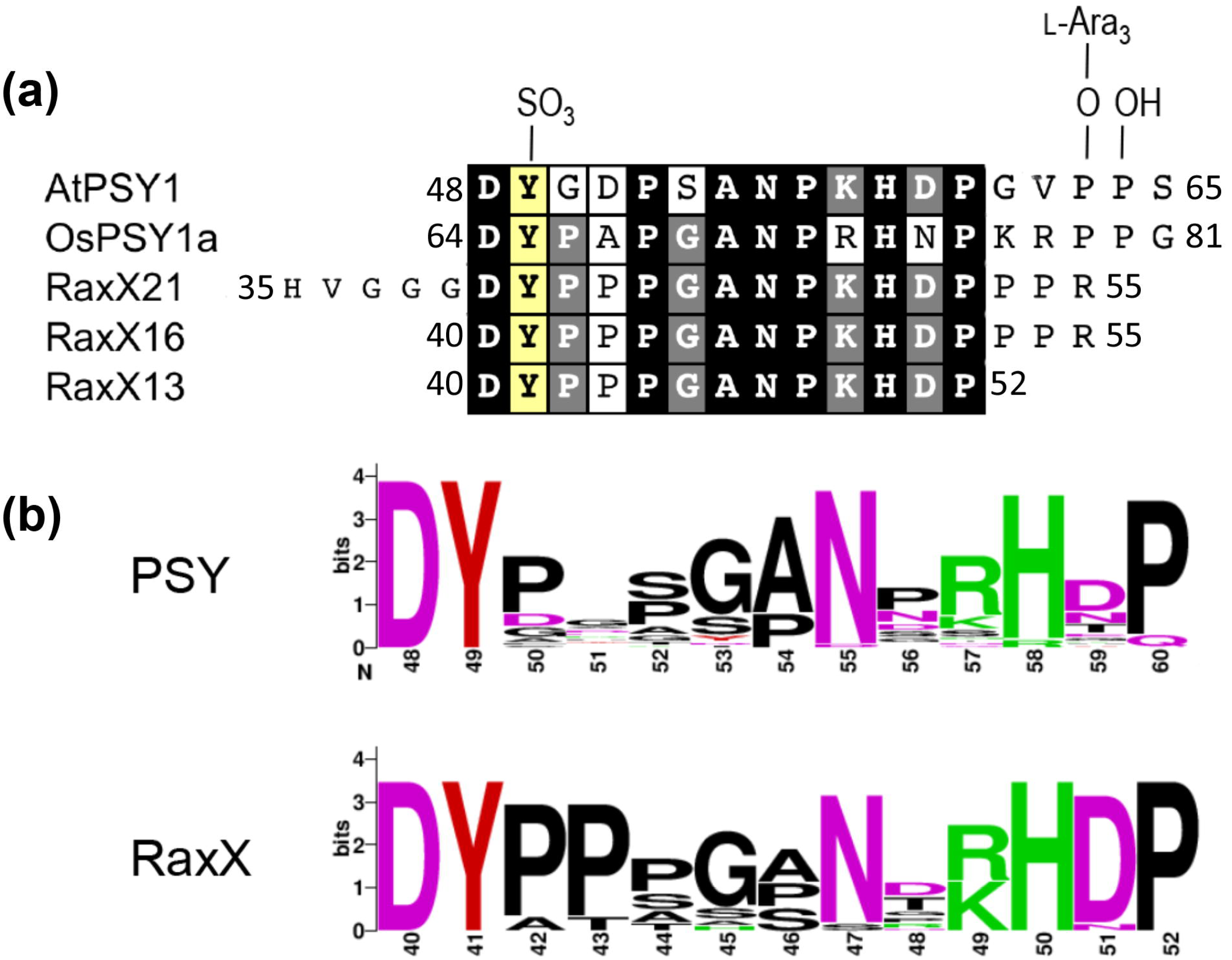
Sequence similarity of RaxX and plant PSYs. a) The mature 18 amino acid AtPSY1 (amino acids 48-65 of the AtPSY1 precursor protein) and a synthetic PSY-like repeat from OsPSY1 (amino acids 64-81 of the AtPSY1 precursor protein) were aligned with the sequences of three synthetic RaxX peptides from *Xoo* strain PXO99. The numbers adjacent to the sequence indicate the amino acid positions of the terminal peptide residues within the predicted precursor protein. Endogenous AtPSY1 has 3 postranslationally modified residues, which are shown at the top of alignment: a sulfotyrosine and two hydroxyprolines. The first hydroxyproline is further modified by chain of three L-arabinose residues (L-Ara_3_). Residues in the black box are identical in all three sequences. The grey boxes indicate a conserved residue in two sequences among AtPSY1, OsPSY1a and RaxX. The sulfated tyrosine is marked in yellow box. b) Sequence logos depicting the amino acid composition in the conserved 13-amino acid region of RaxX and PSY proteins. The logos were generated from 34 PSY orthologs (Fig. S1) and 17 non-redundant RaxX13 sequences (Table S1).

Sequence analysis revealed that RaxX21 is similar to the peptide hormone PSY (*p*lant peptide containing *s*ulfated *t*yrosine), which promotes cellular proliferation and expansion in Arabidopsis (Amano *et al.,* 2007) (Pruitt *et al.,* 2015). Arabidopsis PSY1 (AtPSY1) is the best-characterized member of the plant PSY peptide family. AtPSY1 is an 18-amino acid glycopeptide with a single sulfotyrosine residue (Fig. 1A) (Amano *et al.,* 2007) that is secreted, processed from a 75 amino acid precursor and promotes root elongation primarily through regulation of cell size. AtPSY1 is widely expressed in Arabidopsis tissues (Amano *et al.,* 2007). AtPSY1 promotes acidification of the apoplastic space through activation of membrane proton pumps (Fuglsang *et al.,* 2014). This acidification is thought to activate pH-dependent expansins and cell wall remodeling enzymes that loosen the cellulose network (Cosgrove, 2000; Hager, 2003). Concomitant water uptake by the cell leads to cellular expansion. In addition to PSY, plants produce three other classes of tyrosine sulfated peptides: *p*hyto*s*ulfo*k*ine (PSK) (Matsubayashi & Sakagami, 1996), *r*oot meristem *g*rowth *f*actor (RGF) (Matsuzaki *et al.,* 2010) and Casparian strip integrity factor (CIF) (Doblas *et al.,* 2017; Nakayama *et al.,* 2017). PSK, RGF and CIF are also processed, secreted, and play roles in regulation of growth, development and Casparian strip diffusion barrier formation in the root.

Here we demonstrate that RaxX peptides derived from diverse *Xanthomonas* species promote root growth, mimicking the growth promoting activities of PSY peptides. We also show that a *Xanthomonas* strain lacking *raxX* is impaired in its ability to infect rice lacking XA21, suggesting that RaxX is a virulence factor. Unlike RaxX, PSY peptides do not activate XA21- mediated immunity. Thus, XA21 is a highly selective immune receptor capable of specifically recognizing the bacterial mimic. Based on these findings we propose a model whereby *Xoo* and other *Xanthomonas* strains produce RaxX to reprogram the host environment by hijacking PSY signaling. XA21 later evolved to recognize and respond to specifically to RaxX.

## Materials and Methods

### Identification of Putative RaxX proteins

Putative PSY orthologs were identified by NCBI Protein BLAST analysis using the default settings for short sequences (Altschul *et al.,* 1990). For *Solanum lycopersicum* BLAST was performed using the Sol Genomics Network with the BLOSUM 62 matrix (https://solgenomics.net/tools/blast/). Proteins were identified from a single source for each plant: *Arabidopsis thaliana* Col-0 (refseq_protein, taxid: 3702), *Oryza sativa* Nipponbare (refseq_protein, taxid: 39947), *Triticum aestivum* Chinese Spring (taxid:4565), *Musa acuminata* subsp. *Malaccensis* (refseq_protein, taxid 214687), *S. lycopersicum* cv. Heinz 1706 (ITAG release 2.40). BLAST was initially performed with the 18 amino acid sequence of AtPSY1 (DYGDPSANPKHDPGVPPS). Criteria for selection were as follows: (1) Candidates must match the query with an expect-value ≤ 20 for NCBI Protein BLAST analysis (PAM 30 matrix), (2) Candidates must have an invariant Asp-Tyr at the beginning of the query, (3) The full length protein must be between 60 and 200 amino acids with the PSY-like motif in the second half, (4) The protein must be predicted to have a secretion signal by SignalP 4.0 (Petersen *et al.,* 2011). Additional candidates were identified by subsequent iterative BLAST with the 18 amino acid RaxX sequences from candidate RaxX proteins identified in the initial BLAST. The final list is shown in Figure S1. If multiple splicing variants were identified in the search, only one was listed.

### Sequence analysis and visualization

The sequence alignments in S9 were generated with Geneious software using default parameters (Kearse *et al.,* 2012). Sequence logos (Fig. 1b) were constructed using WebLogo (Schneider & Stephens, 1990; Crooks *et al.,* 2004) with the 13-amino acid RaxX sequences shown in Table S1 and the PSY ortholog sequences in Fig. S1. The bit score for a given residue indicates the conservation at that position, while the size of the individual letters within the stack indicate relative frequency of that amino acid at the position.

### Arabidopsis growth conditions

All *Arabidopsis thaliana* used in this study were in the Col-0 background. The *AtTPST* mutant, *tpst-1,* (SALK_009847) and homozygous *At1g72300* mutant (SALK_072802C) were obtained from the Arabidopsis Biological Resource Center (ARBC). A homozygous *tpst-1* line was isolated from progeny of the SALK_009847 seeds. The *AtPSKR1/AtPSKR2/At1g72300* triple receptor mutant (Mosher *et al.,* 2013) was obtained from Birgit Kemmerling’s laboratory. Plants were grown on the indicated media or on Sungro professional growing mix under continuous light.

### RaxX and PSY1 peptides

The peptides used in this study are listed in Table S2. All peptides other than RaxX21-Y are tyrosine sulfated as indicated (Y^S^). The synthetic AtPSY1 peptide used in these experiments lacks the hydroxy- and L-Ara_3_-modifications at the C-terminus. The natural processed, modified state of OsPSY1a is not known. The 18-amino OsPSY1a acid peptide was synthesized based on alignment with AtPSY1. RaxX13-sY was obtained from Peptide 2.0. All other peptides were obtained from Pacific Immunology. One batch of peptides was tested for each sequence. The peptides were resuspended in ddH_2_O.

### Arabidopsis root growth assays

Arabidopsis seeds were treated with 30% bleach for 12 minutes and then washed 4-5 times with autoclaved water. Sterilized seeds were incubated in the dark at 4°C for 3-4 days. Plates were prepared with 0.5 × Murashige and Skoog (MS) medium with vitamins (Caisson, MSP09), 1% sucrose, pH 5.7, 0.5% Phytagel (Sigma, P8169). Peptide (or water for mock treatments) was added to the indicated concentration (from a 1 mM stock) just before pouring into a plate. Seeds were placed on the plate (20 seeds per plate), and the lids were secured with Micropore surgical tape (1530-0). Plates were incubated vertically under continuous light (55 μmol m^-2^ s^-1^) at 24°C. Seedlings with delayed germination were marked after 3 days, and were not included in the analysis. Root lengths were measured after 8 days.

### Arabidopsis live imaging of root growth

Live imaging of roots was performed as described previously with modifications to the media (Duan *et al.,* 2013; Geng *et al.,* 2013). Sterilized *tpst-1* seeds were grown on 1% agar media containing 1× MS nutrients (Caisson, MSP01), 1% sucrose, and 0.5 g l^-1^ MES, adjusted to pH 5.7 with KOH. After 6 days, seeds were transferred to 0.5% Phytagel (Sigma, P8169) media containing 0.5× MS (Caisson, MSP09, 1% sucrose, and 0.5 g l^-1^ MES, adjusted to pH 5.7 with KOH) with or without the indicated peptides. Imaging and semiautomated image analysis were performed as described previously (Geng *et al.,* 2013).

### Rice root growth assays

Seeds of *Oryza sativa* sp. *japonica* cultivars Kitaake (lacking the *Xa21* gene), a transgenic line of Kitaake carrying *Xa21* (XA21-Kitaake), Taipei 309 (TP309), (lacking the Xa21 gene), or a transgenic line of TP309 carrying *Xa21* driven by its native promoter (XA21-TP309) were dehusked and sterilized with 30% bleach for 30 min. The seeds were washed 4-5 times with water and plated to cups with 50 mL 0.5×MS (Caisson MSP09), 1% sucrose (pH 5.7 with KOH/ NaOH) containing 0.25% Phytagel. Peptides were added to 100 nM just before pouring into the cups. 20 seedlings were added per cup, and the cups were sealed with clear lids. The seedling roots were measured after 4-6 day incubation in a 28°C chamber with 13 h/11 h light/dark cycle and a light intensity of 15 μmol m^-2^ s^-1^.

### ROS Assays

Kitaake and XA21-Kitaake rice plants were grown as previously described (Pruitt *et al.,* 2015). Briefly, seeds were geminated on water-soaked paper and transplanted in sandy soil in 5.5 inch square pots. Plants were grown in tubs filled with fertilizer water in greenhouse. Six weeks after planting the rice was transferred to a growth chamber set to 28 °C/24 °C, 80%/85% humidity, and 14/10 h lighting for the day/night cycle. ROS assays were carried out using leaves of 6-week-old rice plants as described previously (Pruitt *et al.,* 2015). Briefly, leaves were cut longitudinally along the mid vein and then transversely into 1- to 1.5-mm-thick leaf pieces. After overnight incubation floating on sterile water, leaf pieces were transferred into a 96-well white plate (2 pieces per well). Each well contained 100 μl of excitation solution [0.2 mM L-012 (Wako) and 50 μg ml^-1^ horseradish peroxidase (Sigma)]. The indicated concentration of peptides was added (or water for mock control), and chemiluminescence was measured for 90 minutes with a TriStar (Berthold) plate reader.

### Xanthomonas inoculation on rice

TP309 and XA21-TP309 were greenhouse grown as described above for Kitaake. Plants were inoculated 3 days after transfer using the scissors clipping method (Kauffman *et al.,* 1973). PXO99 strains were grown on peptone sucrose agar (PSA) plates at 28 °C with the appropriate antibiotic(s). The bacteria were resuspended in water at a density of 10^6^ colony forming units per mL. Water soaked lesions were measured 14 days after inoculation. Bacterial growth analysis *in planta* was performed as previously described (Bahar *et al.,* 2014). PXO99 strains used in this study were previously reported (Pruitt *et al.,* 2015). PXO99Δ*raxX* is a marker free mutant and PXO99Δ*raxST* is a marker exchange mutant with a spectinomycin resistance gene. The *raxX* and *raxST* sequences including their predicted promoter were cloned into pVSP61 vector (Loper & Lindow, 1994) and transformed into PXO99 strains.

## Results

### RaxX is similar in sequence to PSY peptides

The region of similarity between RaxX from *Xoo* and AtPSY1 corresponds to amino acids 40-52 of RaxX. RaxX and AtPSY1 share 10 identical residues over this region (Fig. 1a). RaxX is sulfated by the bacterial sulfotransferase RaxST on Y41, which corresponds to the sulfated residue of AtPSY1 (Amano *et al.,* 2007; Pruitt *et al.,* 2015). An aspartate precedes the sulfated tyrosine in both RaxX and AtPSY1. The presence of a nearby acidic residue is a common hallmark of tyrosine sulfation sites (Moore, 2009).

We extended our analysis to include PSY orthologs and RaxX peptides from diverse species (Fig. S1-2, Table S1). BLAST search using the18 amino acid AtPSY1 as a query identified 8 PSY-like proteins in rice (Fig. S1). One of the rice PSY proteins, OsPSY1 (Os05g40850), has four nearly identical PSY-like repeats, the first of which (OsPSY1a) is shown in Fig 1. Analysis of Arabidopsis using the same criteria also revealed a total of eight PSY-like proteins including the three that had been previously identified (Fig. S1) (Amano *et al.,* 2007; Matsubayashi, 2014). We also identified PSY-like proteins in tomato, banana and wheat, three diverse and economically important crops (Fig. S1). Alignment of PSY peptides from these different species revealed a highly conserved 13-amino acid region beginning with the aspartate-tyrosine residue pair (Fig. S1). This 13-amino acid sequence corresponds precisely to the region of sequence similarity between RaxX and AtPSY1 (Fig. 1a).

Alignment of the RaxX sequences from diverse strains reveals a region of high conservation immediately around the tyrosine, which is sulfated in *Xoo* strain PXO99 (Fig. S2). Sequence logos were constructed for the PSY-like motif using the identified RaxX and PSY sequences (Fig. 1b). These logos further highlight the similarity of 13-amino acid region of RaxX and PSY sequences. Residues that are highly variable in RaxX are also highly variable in PSY. Based on the similarity of RaxX and PSY peptides and the finding that RaxX is also tyrosine sulfated (Pruitt *et al.,* 2015), we hypothesized that RaxX serves as a functional mimic of PSY peptides and that RaxX may have PSY-like activity.

### RaxX promotes root growth similar to PSY peptides

To test our hypothesis that RaxX is a functional mimic of PSY peptides, we evaluated the effect of RaxX21 treatment on root growth. We first tested the peptides on Arabidopsis seedlings, because PSY signaling has been studied exclusively in this system. RaxX21-sY promoted root growth in a similar manner to that observed for AtPSY1 in Arabidopsis (Fig. 2a, b). After 8 days on media containing 100 nM RaxX21-sY, the average root length of Col-0 seedlings was 61 mm whereas seedlings grown on plates without peptide had an average root length of 54 mm. Similar root growth-promoting effects were observed in experiments using AtPSY1 and OsPSY1a peptides (Fig. 2a, b).

**Figure 2.**
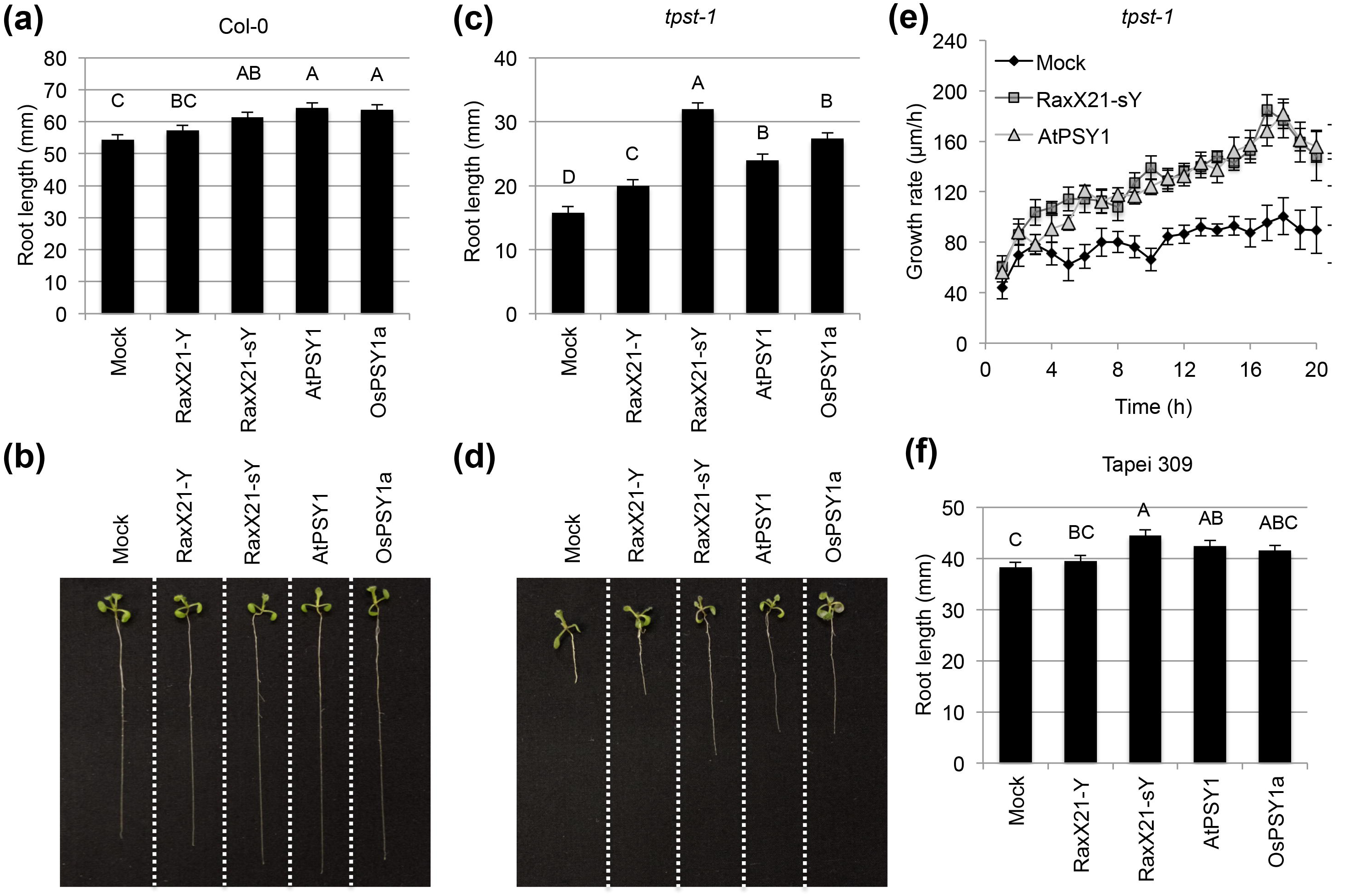
Sulfated RaxX21 promotes root growth in *Arabidopsis* and rice. Root lengths of Arabidopsis a) Col-0 or c) *tpst-1* seedlings grown on 0.5× MS vertical plates with or without 100 nM of the indicated peptides. Bars indicate the average seedling root length measured after 8 days (n ≥ 18). b and d) 8-day old Col-0 and *tpst-1* seedlings grown as in 2a and c, respectively. e) Growth rate of six day old *tpst-1* seedlings following transfer to 0.5× MS plates containing 250 nM RaxX21-sY, 250 nM AtPSY1, or lacking peptide (Mock) (n ≥ 7). Growth was monitored by continual imaging over 20 h. f) Root lengths of 6-day old rice seedlings (Tapei 309) grown on 0.5× MS with or without 100 nM of the indicated peptides (n ≥ 37). Error bars indicate standard error. Statistical analysis was performed using the Tukey-Kramer honestly significant difference test for mean comparison using the JMP software. Different letters represent significant differences within each plant genotype (p ≤ 0.05).

We also performed root growth experiments on an Arabidopsis line lacking AtTPST, the tyrosine sulfotransferase responsible for modification of PSY, PSK and RGF peptides (Komori *et al.,* 2009; Matsuzaki *et al.,* 2010). *tpst-1* mutant plants are dwarf and have stunted roots (Komori *et al.,* 2009). Because this mutant lacks endogenous PSY, PSK and RGF signaling, effects of exogenous application of sulfated peptides can be better quantified (Igarashi *et al.,* 2012; Mosher *et al.,* 2013). Consistent with earlier reports, we observed that mock treated *tpst-1* mutant seedlings have much shorter roots than Col-0 (Fig. 2a-d). Treatment of *tpst-1* plants with RaxX21-sY or AtPSY1 increases root growth 1.5-2 fold relative to mock treatment (Fig 2c, d).

We determined the minimum concentration of RaxX21-sY needed to induce root growth in Arabidopsis. *tpst-1* seeds were grown on plates containing 0.1-250 nM peptide. RaxX21-sY was effective at inducing root growth at concentrations in the low nanomolar range (Fig. S3). This activity is comparable to PSK (Fig. S3). Nonsulfated RaxX21 (RaxX21-Y) also promoted root growth, but was less active than the sulfated version (Fig. 2a-d, S3). AtPSY1 was less active than RaxX21-sY and PSK. We hypothesize that the reduced potency of the synthetic AtPSY1 used in this study was due to the lack of glycosylation (See materials and methods). Glycosylation of AtPSY1 was previously shown to be important for full activity (Amano *et al.,* 2007).

We next used a live root imaging system (Duan *et al.,* 2013; Geng *et al.,* 2013) to assess changes in root growth rate upon exposure to RaxX21-sY. Root growth of *tpst-1* seedlings on plates containing 250 nM RaxX21-sY, AtPSY1 or no peptide (Mock) was monitored over 24 h. Within 4-5 hours, seedlings grown on RaxX21-sY- or AtPSY1-containing plates had an increased root growth rate compared to seedlings on mock plates (Fig. 2e).

Because RaxX21-sY comes from the rice pathogen *Xoo,* we tested whether this peptide also has growth promoting activity in rice seedlings. AtPSY1 and RaxX21-sY treatment significantly enhanced root growth on rice varieties Tapei 309 (Fig. 2f) and Kitaake (Fig. S4). We also tested if the root growth promotion activity is attenuated in the presence of XA21. We found that treatment of RaxX21-sY still induced longer roots in XA21-TP309 plants (Fig. S5). We hypothesize that RaxX21-sY fails to activate XA21 in young seedlings, because XA21- mediated immune response is developmentally controlled in rice (Century *et al.,* 1999). Collectively, these results indicate that RaxX21-sY promotes root growth in a similar manner to PSY and PSK peptides in both Arabidopsis and rice.

### RaxX induces root growth through the same signaling pathway as PSY1

To determine if RaxX induces root growth using the same signaling pathway as AtPSY1, we grew Arabidopsis seedlings on plates containing both RaxX and AtPSY1 peptides. Roots of Arabidopsis seedlings grown on plates containing 100 nM RaxX21-sY and 100 nM AtPSY1 were similar in length to those grown on plates with 100 nM RaxX21-sY alone (Fig. 3). Similar results were observed when seedlings were co-treated with 100 nM RaxX21-sY and 100 nM PSK (Fig. 3). The observation that RaxX, AtPSY1, and PSK do not have additive effects on root growth suggests that these peptides induce root growth via the same pathway. Alternatively, it may be that the 100 nM RaxX21-sY treatment already reached the maximum growth potential (Matsuzaki *et al.,* 2010).

**Figure 3.**
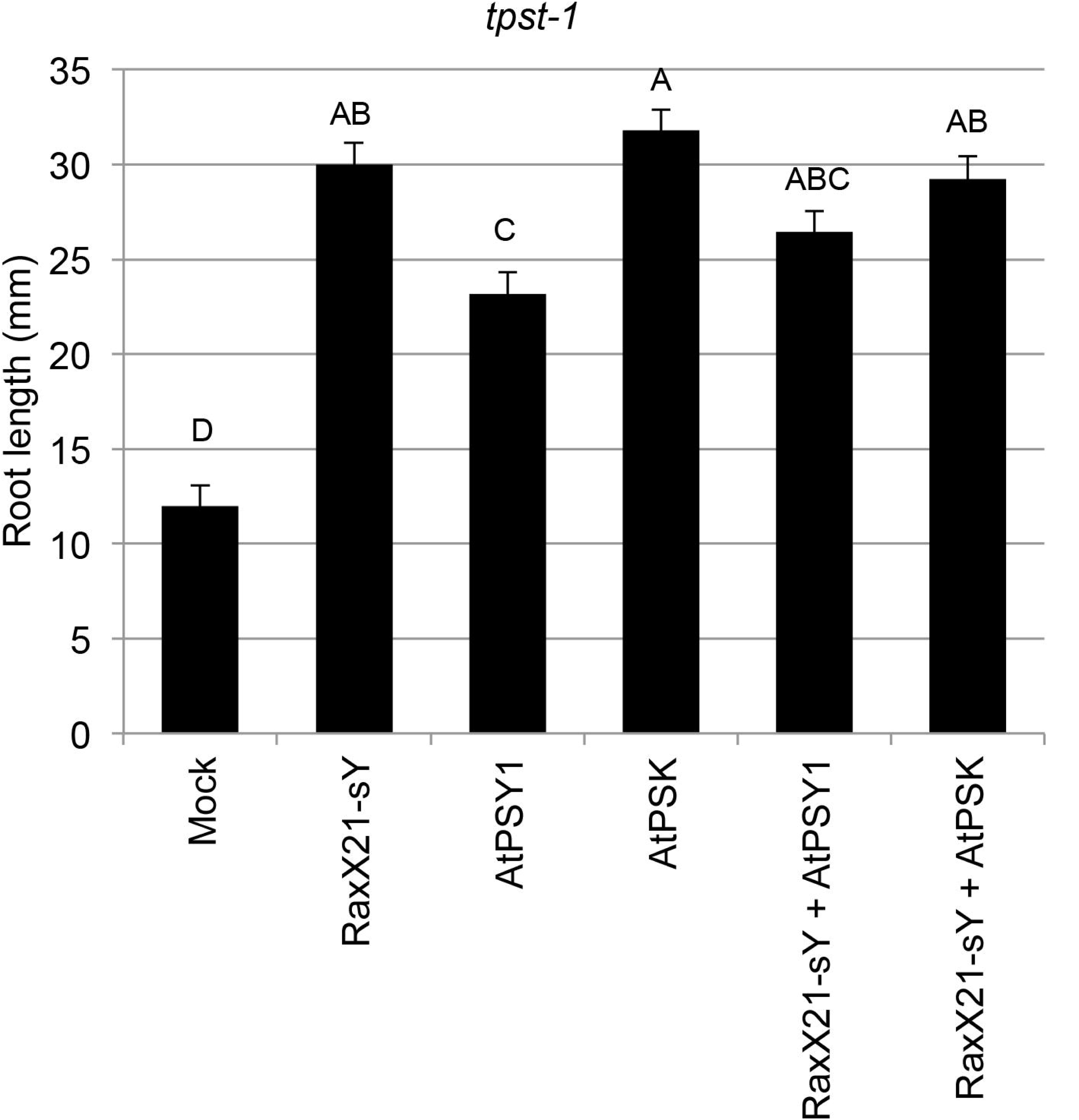
RaxX, AtPSY1, and PSK do not have additive effects on root growth in Arabidopsis. *tpst-1* seedlings were grown on 0.5× MS vertical plates with or without 100 nM of each of the indicated peptides. Bars indicate the average seedling root length measured 8 days after plating seeds (n ≥ 18). Error bars indicate standard error. Statistical analysis was performed using the Tukey-Kramer honestly significant difference test for mean comparison using the JMP software. Different letters represent significant differences within each plant genotype (p ≤ 0.05). Experiments were performed at least two times with similar results.

### At1g72300 is not required for induction of root growth by RaxX or AtPSY1

The leucine-rich repeat receptor kinase encoded by *At1g72300* has been proposed to serve as the AtPSY1 receptor (Amano *et al.,* 2007). We therefore tested if At1g72300 is required for perception of RaxX21-sY. For these assays we used the *At1g72300* mutant line SALK_072802C. This is the same line used in all published studies of PSY1/At1g72300, (Amano *et al.,* 2007; Mosher & Kemmerling, 2013; Mosher *et al.,* 2013; Fuglsang *et al.,* 2014; Mahmood *et al.,* 2014), and was shown to have the lowest transcript level of available mutants (Fuglsang *et al.,* 2014). We independently validated the mutant genotype (Fig. S6). We found that treatment of the *At1g72300* mutant line with either RaxX21-sY or AtPSY1 increased root growth in a similar manner to that observed for treatment of wild type Col-0 seedlings (Fig. 2a, b, 4). We also found that a mutant lacking At1g72300 and the homologous PSK receptors, AtPSKR1 and AtPSKR2, (*pskr1/pskr2/At1g72300*) also responds to RaxX and AtPSY1 treatment (Fig. 4). *pskr1/pskr2/At1g72300* did not respond to synthesized Arabidopsis PSK (AtPSK), whereas PSK promotes root growth of wild-type Col-0 and *At1g72300* (Fig. 4). These results indicate that At1g72300 is not required for perception of RaxX21-sY or AtPSY1.

**Figure 4.**
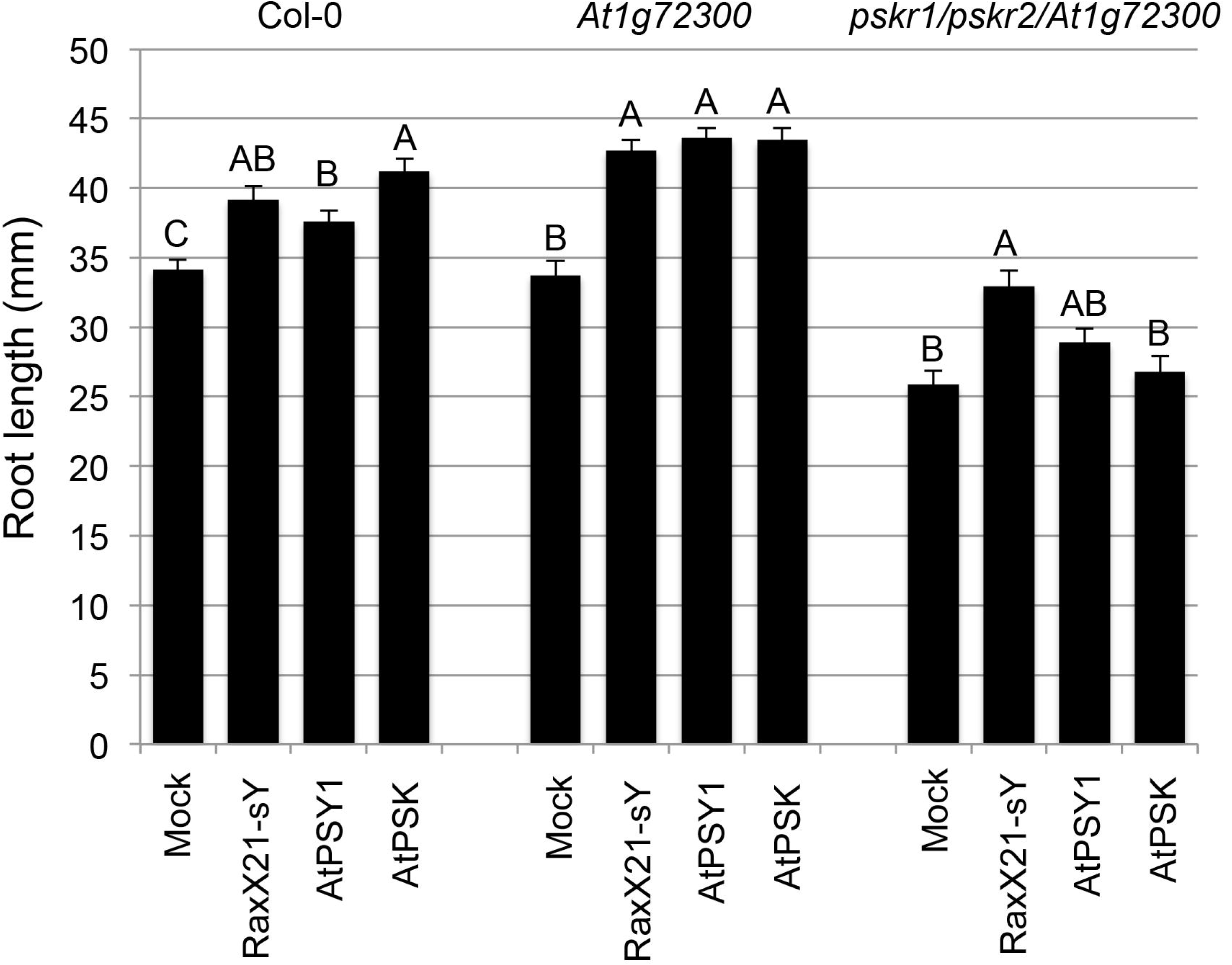
The Arabidopsis gene *At1g72300* is not required for RaxX- and PSY-induced root growth. Arabidopsis Col-0, *At1g72300* or *AtPSKR1/AtPSKR2/At1g72300* triple receptor mutant seeds were grown on 0.5× MS plates with or without 100 nM of the indicated peptides. Root lengths were measured 8 days after placing seeds on plates. Error bars indicate standard error (n ≥ 22). Statistical analysis was performed using the Tukey-Kramer honestly significant difference test for mean comparison using the JMP software. Different letters represent significant differences within each plant genotype (p ≤ 0.05). The experiment was performed at least three times with similar results.

### RaxX21-sY and PSY do not attenuate elf18-induced growth inhibition

Exogenous addition of PSK has previously been shown to attenuate the Arabidopsis immune response to biotrophic pathogens (Igarashi *et al.,* 2012; Mosher & Kemmerling, 2013; Mosher *et al.,* 2013). Although PSK and AtPSY1 share no sequence similarity, they have nevertheless been hypothesized to serve similar roles (Mosher & Kemmerling, 2013; Mosher *et al.,* 2013; Matsubayashi, 2014). Thus, we hypothesized that induction of PSY signaling by PSY or RaxX21-sY may also attenuate plant immune responses. To test this hypothesis, we employed a seedling growth inhibition assay. Arabidopsis seedlings were grown in the presence of the bacterial elicitor elf18, which causes activation of immune response and impairs growth. We demonstrated that co-incubation of seedlings with PSK attenuates elf18-mediated growth inhibition as previously reported (Igarashi *et al.,* 2012) (Fig. S7). However, RaxX21-sY and AtPSY1 do not prevent elf18-triggered growth inhibition in Arabidopsis under the conditions tested (Fig. S7). These results indicate that RaxX21-sY and PSY1 do not have the same effects on immune modulation as PSK in Arabidopsis seedlings in response to elf18 treatment.

### RaxX and PSY peptides differentially activate PSY-like growth promotion and XA21-immune responses

Activation of XA21-mediated immunity by RaxX21-sY triggers a number of immune responses including production of reactive oxygen species (ROS), induction of marker gene expression, and production of ethylene (Pruitt *et al.,* 2015). These immune responses are tightly regulated, because aberrant activation of immunity can have negative effects on plant growth and health (Spoel & Dong, 2012; Rodriguez *et al.,* 2015). We therefore hypothesized that XA21 would specifically recognize RaxX but not the homologous PSY peptides.

We have previously shown that RaxX21-sY treatment induces robust ROS production in rice leaves expressing XA21 (Pruitt *et al.,* 2015). Therefore, to assess XA21-mediated recognition of the sulfated peptides, we measured ROS production in XA21 rice leaves upon treatment with water, RaxX21-sY, AtPSY1, or OsPSY1a (Fig. 5a). Unlike RaxX21-sY, AtPSY1 and OsPSY1a failed to induce ROS production in XA21 rice leaves. Robust ROS production was not observed in rice leaves lacking XA21 (Fig. 5b). PSK also failed to activate XA21-mediated immune response (Fig. 5a, b). These results suggest that the XA21 and PSY receptor(s) have different specificities. PSY signaling with respect to primary root growth is activated by both PSY and RaxX (Fig. 2), whereas the XA21-mediated immune response is only activated by RaxX.

**Figure 5.**
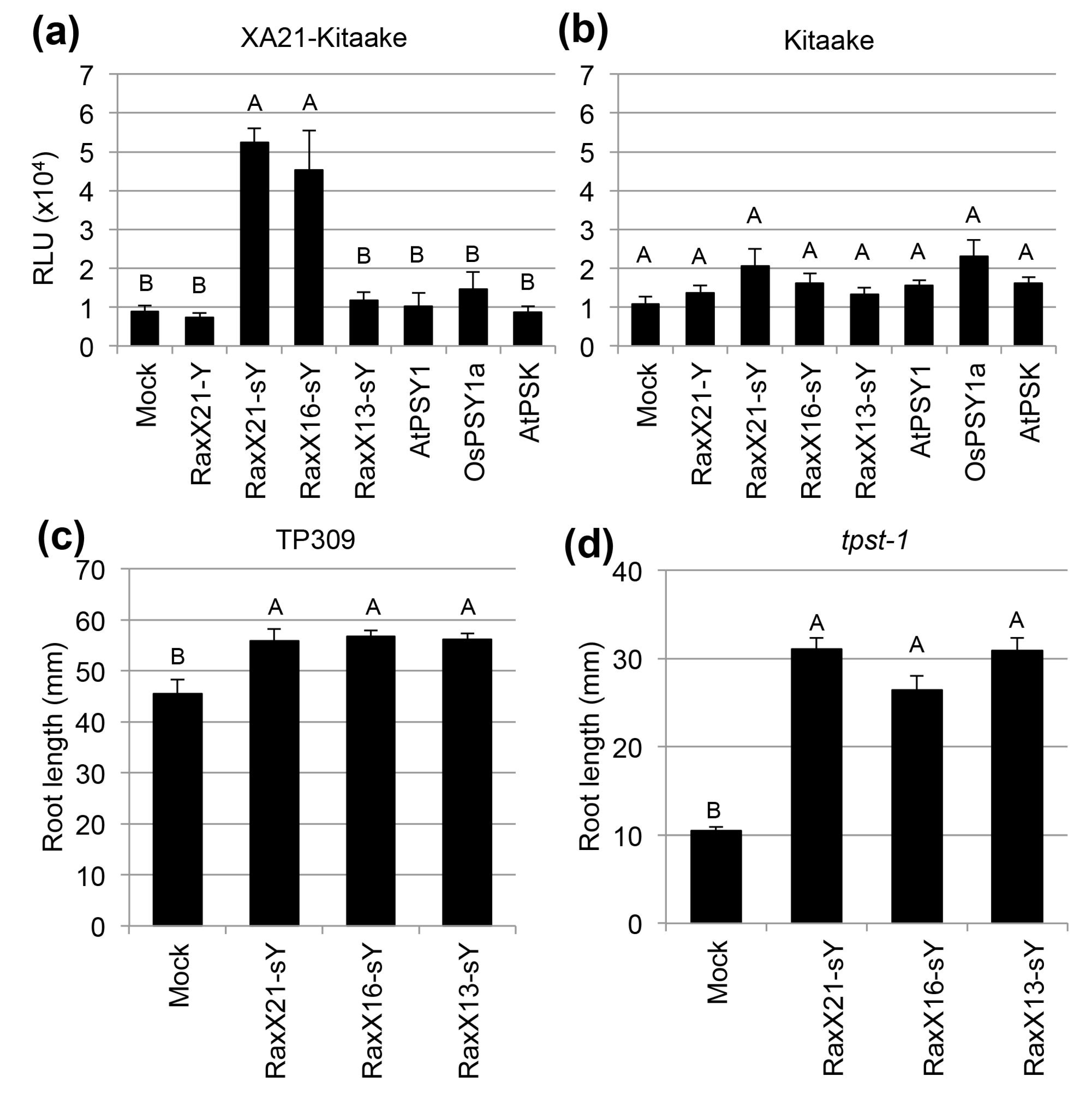
Differential activities of PSY and RaxX peptides in growth promotion and activation of XA21-mediated immunity. ROS production in leaves of a) XA21 rice (XA21- Kitaake) and b) wild type rice (Kitaake) treated with H_2_O (Mock) or 500 nM of the indicated peptide. Bars represent average ROS production over 90 min following addition of peptide (n = 6). RLU stands for relative light units. c) TP309 seeds were grown on 0.5× MS media for with or without 100 nM of the indicated peptides. Root lengths were measured 5 days after placing seeds on plates (n ≥ 25). d) Arabidopsis *tpst-1* seeds were grown on 0.5× MS vertical plates with or without 100 nM of the indicated peptides. Root lengths were measured 8 days after placing seeds on plates (n ≥ 16). Error bars indicate standard error. Statistical analysis was performed using the Tukey-Kramer honestly significant difference test for mean comparison using the JMP software. Different letters represent significant differences within each plant genotype (p ≤ 0.05). Experiments were performed at least two times with similar results.

To further delineate the region of RaxX required for PSY-like activity and activation of XA21, we synthesized two smaller RaxX peptides based on similarity to AtPSY1. RaxX16-sY begins with the aspartate (D40) at the beginning of the PSY-like motif (Fig. 1a). RaxX13-sY also begins with D40 but is C-terminally truncated relative to RaxX21-sY and RaxX16-sY (Fig. 1a). RaxX13-sY contains the region of highest similarity shared between the RaxX and PSY peptides (Fig. 1, S1). Both the RaxX13-sY and RaxX16-sY peptides are still capable of promoting root growth in Arabidopsis and rice (Fig. 5c, d). We next tested whether these peptides could activate XA21-mediated immunity in the same manner as RaxX21-sY (Pruitt *et al.,* 2015). For this purpose, ROS production was measured in detached XA21 rice leaves treated with water, RaxX13-sY, RaxX16-sY, or RaxX21-sY. RaxX16-sY and RaxX21-sY triggered a ROS response characteristic of the XA21-mediated immune response. In contrast, treatment with RaxX13-sY did not induce ROS production in XA21 rice leaves (Fig. 5a). Thus, RaxX13-sY is able to induce AtPSY1-like growth effects, but fails to activate an XA21-mediated immune response. These experiments reveal that RaxX residues 53-55, which are present in RaxX16 but not RaxX13, are important for activation of XA21 but are not required for root growth promoting activity.

### RaxX from diverse Xanthomonas species have PSY activity

We next asked whether RaxX from other *Xanthomonas* strains also have PSY-like activity. To address this question, we synthesized 24-amino acid peptides covering the PSY-like region for three different RaxX sequences from *X. oryzae* pv. *oryzicola* strain BSL256 (RaxX24-Xoc-sY), *X. campestris* pv *musacearum* strain NCPPB4394 (RaxX24-Xcm-sY), and *X. euvesicatoria* strain 85-10 (RaxX24-Xe-sY) (Table S2). *Xoc, Xcm,* and *Xe* are pathogens of rice, banana, and tomato/pepper, respectively (Table S1). *Xoc* colonizes the mesophyll of rice, whereas *Xoo* colonizes the xylem. All three RaxX sulfated peptides promoted root growth on Arabidopsis seedlings in a manner similar to that of RaxX21-sY derived from *Xoo* strain PXO99 (Fig. 6). In other words, the proteins encoded by diverse allelic variants of *raxX* retain PSY like activity. These results demonstrate that the use of RaxX as a mimic of plant PSYs is employed by many *Xanthomonas* species that infect diverse plant species.

**Figure 6.**
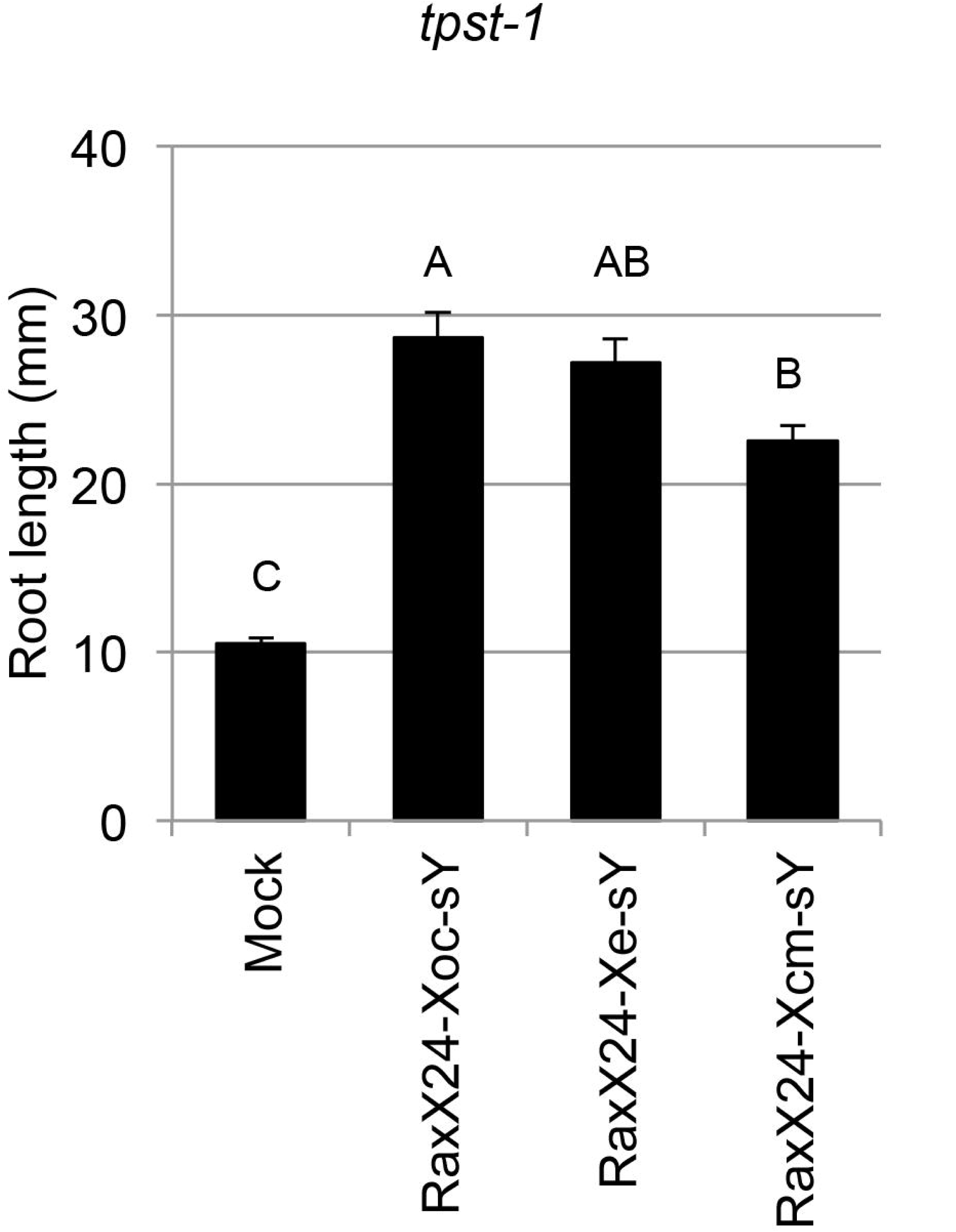
RaxX peptides derived from RaxX encoded by *Xoc, Xe,* and *Xcm* promote root growth in Arabidopsis seedlings. *tpst-1* seedlings were grown on 0.5× MS vertical plates with or without 100 nM of the indicated peptides. Bars indicate the average seedling root length measured after 8 days (n ≥ 18). Error bars indicate standard error. Statistical analysis was performed using the Tukey-Kramer honestly significant difference test for mean comparison using the JMP software. Different letters represent significant differences within each plant genotype (p ≤ 0.05). Experiments were performed at least two times with similar results.

### RaxX facilitates Xoo infection

In some cases, the ability of a pathogen to mimic a host biological process can facilitate pathogen infection (Weiler *et al.,* 1994; Melotto *et al.,* 2006; Mitchum *et al.,* 2012; Chen *et al.,* 2015). We therefore tested if RaxX contributes to the virulence of *Xoo* in plants lacking XA21. We did not observe an effect of RaxX on disease lesion development in TP309 rice leaves using standard scissor clipping inoculation (a high inoculum concentration of 10^8^ colony forming units per mL) (da Silva *et al.,* 2004; Pruitt *et al.,* 2015). Inoculating with a low inoculum concentration is known to reveal subtle virulence differences between strains (Starkey & Rahme, 2009). Thus, we challenged TP309 leaves with PXO99 strains at a density of 10^6^ colony forming units per mL. Under this condition, the PXO99Δ*raxX* strain, but not the complemented strain (*PXO99*Δ*raxX*(*praxX*)), formed shorter lesions compared with wild-type PXO99 (Fig. 7a). We also tested if RaxST-mediated sulfation is required for the virulence activity of RaxX. A PXO99 strain lacking RaxST (PXO99Δ*raxST)* also formed shorter lesion than PXO99 on TP309 rice leaves in low inoculum concentration experiments (Fig. 7a). PXO99*ΔraxST (praxST)* regained the ability to form long lesions similar to the wild-type strain (Fig. 7a). PXO99 wild-type, PXO99Δ*raxX*(*praxX*) and PXO99Δ*raxST*(*praxST*) form short lesions on XA21-TP309 at a lower inoculum concentration suggesting activation of the XA21 immune response (Fig. 7b). As expected, *PXO99*Δ*raxX* and PXO99Δ*raxST* evade XA21-mediated immune response and form longer lesions (Fig. 7b). The bacterial populations of *PXO99*Δ*raxX* and PXO99 Δ*raxST* were less than those of strains PXO99, PXO99Δ*raxX*(*praxX*), and PXO99Δ*raxST* (*praxST*) 12 days after inoculation (Fig. S8). These results suggest that RaxX is a virulence factor that facilitates *Xoo* infection and that RaxST-mediated sulfation is also required for this virulence activity.

**Figure 7.**
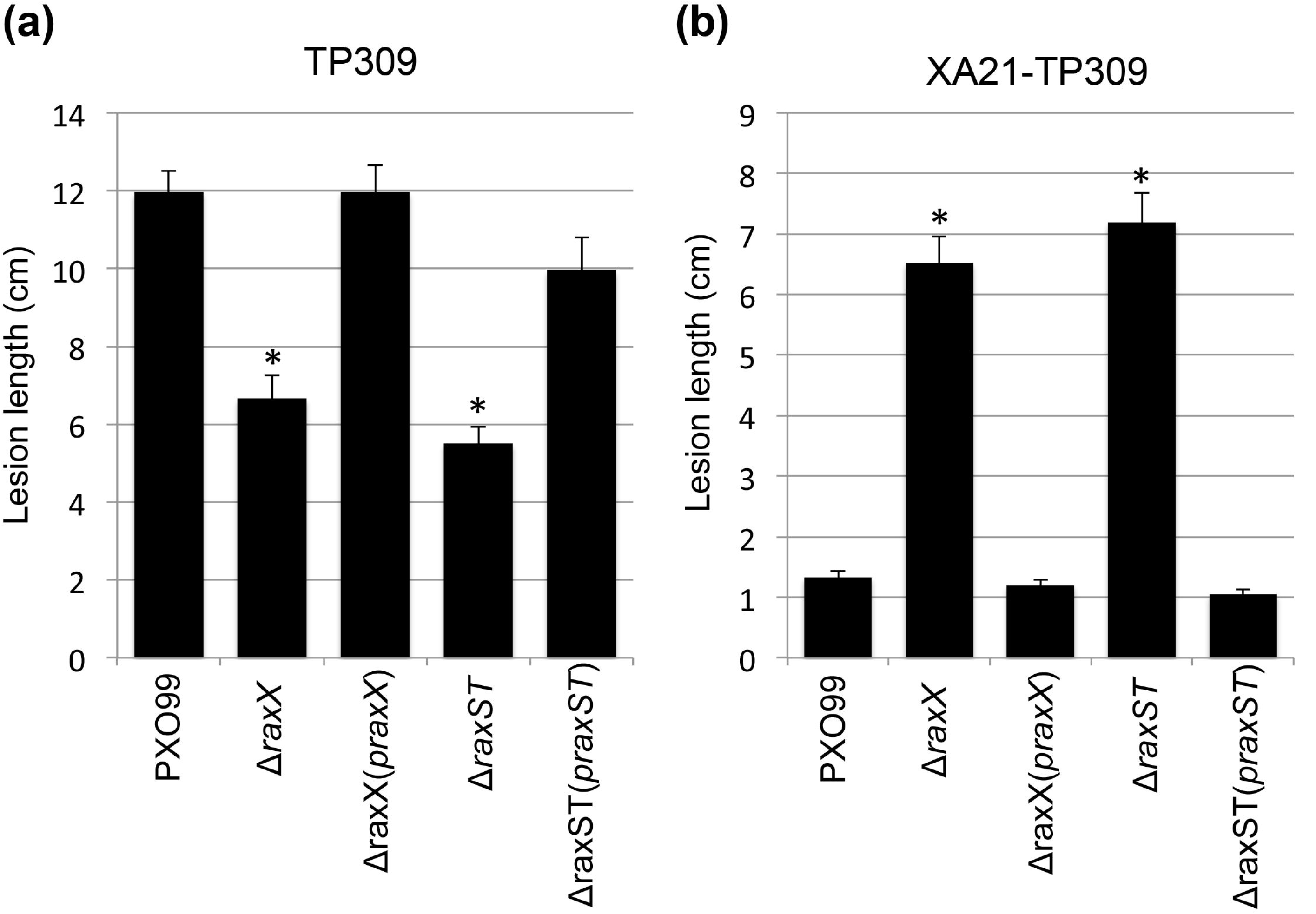
The *Xoo raxX* mutant is impaired in virulence on rice. TP309 (A) and XA21- TP309 (B) were inoculated by clipping with scissors dipped in the indicated *Xoo* suspensions at a density of 10^6^ colony forming units (CFU) per mL. Bars indicate the mean lesion length ± standard error (SE) measured 14 days after inoculation (n ≥24). The ‘*’ indicates statistically significant difference from PXO99 within each plant genotype using Dunnett’s test (α=0.01). Experiments were performed at least five times with similar results.

## Discussion

In a classical evolutionary arms race, both the pathogen and host develop and deploy an arsenal of strategies to infect or resist their partner. For example, many pathogens secrete an array of molecular factors designed to manipulate host biology and suppress the immune response. In turn, plants have developed a set of immune receptors that recognize these molecules or their activities and launch mechanisms to destroy the pathogen, which the pathogen then tries to counter.

The findings reported here and in previous studies, suggest a model where *Xoo* produces, sulfates, and secretes a peptide that mimics PSY peptides (da Silva *et al.,* 2004; Pruitt *et al.,* 2015) (Fig. 8). Plants evolved the receptor XA21 to specifically recognize the bacterial mimic, allowing it to launch a defense response in the presence of the pathogen but not in the presence of the highly similar PSY peptide hormones, which are predicted to be necessary for normal growth and development.

**Figure 8.**
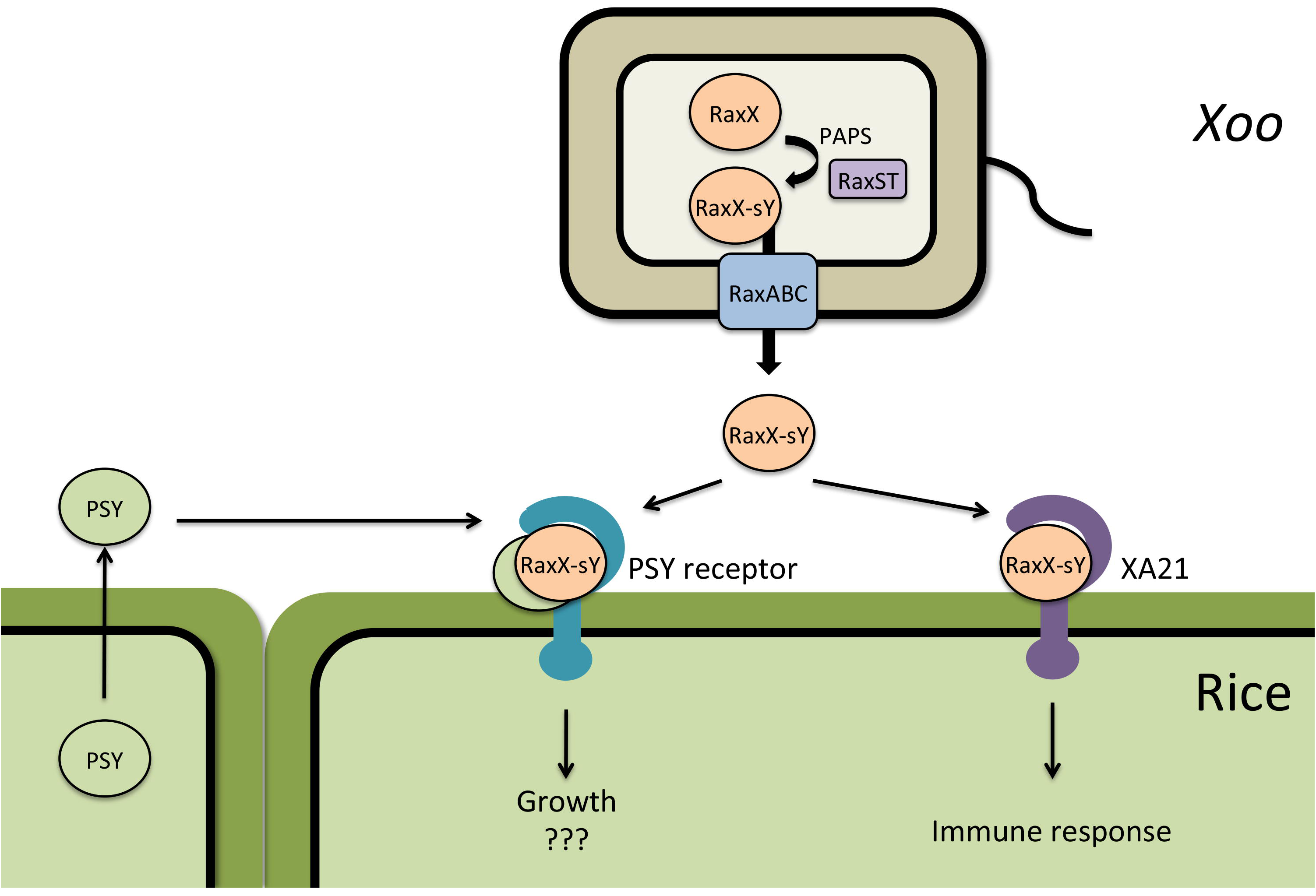
Proposed model of RaxX production and activation of PSY and XA21 signaling. PSY is produced and detected by plant cells to regulate growth. RaxX is produced in *Xoo,* sulfated by RaxST, and secreted by a type I secretion system composed of RaxA, RaxB, and RaxC. Secreted sulfated RaxX induces signaling through the endogenous PSY receptor(s). The wild rice *O. longistaminata* subsequently evolved the immune receptor XA21 which is activated by RaxX, but not endogenous PSY peptides.

The hypothesis that RaxX is a mimic of PSY is well supported by the high level of sequence similarity (Fig. 1), the tyrosine sulfation status of RaxX and PSY peptides (Amano *et al.,* 2007; Pruitt *et al.,* 2015), and the similar growth promoting activities of both peptides (Fig. 2, S3-5). Significantly, both RaxX and PSY1 require tyrosine sulfation for full activity. Tyrosine sulfation is an important posttranslational modification that mediates protein-protein interactions. Plants and animals employ tyrosine-sulfated proteins, to regulate growth, development, immunity and other biological processes. Tyrosine sulfated proteins in animal cells have roles in coagulation, leukocyte adhesion, HIV entry, and chemokine signaling (Farzan *et al.,* 1999; Moore, 2009; Stone *et al.,* 2009).

Due to the similar sequence and functional mimicry in root growth promotion we hypothesize that PSY1 and RaxX target a common cognate plant receptor. The leucine-rich repeat receptor kinase At1g72300 was originally hypothesized to serve as the receptor for AtPSY1 based on the observation that the root length was not increased by exogenous AtPSY1 treatment in an At1g72300 mutant (Amano *et al.,* 2007). However, the At1g72300 mutant line still partially responds to AtPSY1 treatment in proton efflux experiments (Fuglsang *et al.,* 2014), and transcriptomics analysis reveals that many AtPSY1-regulated genes are regulated independently of At1g72300 (Mahmood *et al.,* 2014). We found that RaxX and AtPSY1 still promote root growth in the absence of At1g72300. Collectively, these findings indicate that At1g72300 is not the receptor for PSY peptides or that it is not the only receptor. Additional work is required to understand how PSY and RaxX are perceived in plants.

The precise role of RaxX in *Xoo* biology is not known. Because bacteria have been demonstrated to employ bio-mimics to hijack the plants’ endogenous systems and reprogram the host environment to facilitate pathogen infection (Weiler *et al.,* 1994; Melotto *et al.,* 2006; Mitchum *et al.,* 2012; Chen *et al.,* 2015), we hypothesize that *Xoo* may use RaxX in a similar manner. Here we show that RaxX is required for the full virulence of *Xoo* to infect rice leaves (Fig. 7). *Xoo* is a biotrophic pathogen and thus requires living host tissues, which ensures prolonged supply of carbon and other nutrients necessary for bacterial survival. The ability of *Xoo* to promote the host growth would thus benefit a biotroph (Nino-Liu *et al.,* 2006; Fatima & Senthil-Kumar, 2015).

Xanthomonads enter through hydathodes, natural openings in the leaf, or wounds and multiply in the xylem or mesophyll tissues. To date, growth promoting activities for RaxX or PSY1 have only been demonstrated on roots. We used induction of root growth as an indicator of PSY-like activity in this study because this is a robust well-characterized effect of AtPSY1. It is known, however, that *AtPSY1* is widely expressed in various plant tissues (Amano *et al.,* 2007). Arabidopsis seedlings overexpressing *AtPSY1* not only have longer roots, but also larger cotyledons (Amano *et al.,* 2007). Recently, a PSY-like peptide in soybean is shown to be translocated from the roots to the xylem (Okamoto *et al.,* 2015). These findings suggest that PSY peptides likely have important unidentified roles outside of the roots.

The growth promoting properties of RaxX are reminiscent of the hypertrophy in tomato and pepper leaves induced by the *Xe* effector AvrBs3. AvrBS3 enhances transcription of host genes including auxin-induced and expansin-like genes that contribute to host cell enlargement (Marois *et al.,* 2002). This phenotype is thought to facilitate dissemination because the bacteria are able to multiply in the enlarged cells and escape from the infected site to other plants (Marois *et al.,* 2002; Kay *et al.,* 2007). The AvrBs3 example suggests a possible role for RaxX in bacterial maintenance, persistence or transmission.

In this paper we demonstrate that XA21 can be activated by RaxX16 but not by RaxX13, indicating that the C-terminal end of the RaxX16 sequence (RaxX amino acids 53-55) is required for XA21 recognition. This result may explain why PSY1 cannot activate XA21: PSY1 has C-terminal residues which differ from RaxX16. Residues within the RaxX13 region are also important for recognition by XA21. In a previous study, we identified three residues (44, 46, and 48) of RaxX from *Xoo* that are involved in XA21 activation (Pruitt *et al.,* 2015). Mutation of RaxX P44 and P48 completely abolishes the immunogenic activity of RaxX on XA21-rice. Mutation of A46 has a partial effect. Interestingly, these residues are not required for root growth promoting activity. For example, RaxX24-Xoc contains amino acid differences at positions 44, 46 and 48, but is still capable of inducing root growth in Arabidopsis (Fig. 6, S2, Table S1).

Comparison of the RaxX-Xoo and *RaxX-Xoc* sequences with rice PSY sequences suggests the possibility that RaxX from the *Xanthomonas* strains have evolved to mimic different PSY peptides. The three residues from RaxX-Xoo (strain PXO99) which are required for recognition by XA21 are identical to those in OsPSY1a (Fig. S9). In contrast, the amino acids of RaxX-Xoc (strain BSL256) are similar to those in OsPSY2. If these two peptides have evolved to mimic different PSY peptides, it would indicate that there are multiple PSY receptors in rice, which differentially recognize diverse PSY peptides. Multiple receptors have been reported for RGF peptides. It is not yet clear if the RGF receptors have different affinities for specific RGF peptides (Shinohara *et al.,* 2016). Using multiple receptors and multiple ligands with different affinities would allow for a more complex and tunable signaling network.

To further investigate the possibility that RaxX may have evolved to mimic specific host PSY peptides, we compared the sequences of RaxX13 and PSY from various species (Fig. 1B, S10). We did not observe a correlation between the sequences of RaxX from the pathogen and PSYs from a compatible host (Fig. S10). However, alignment of the 13-amnio acid region did highlight variation at positions 5, 7, and 9. These residues correspond to RaxX amino acids 44, 46, and 48, which are important for XA21 recognition. Notably, the variation is not random. For example, the most common amino acids in position 5 of the sequences analyzed are serine and proline in both RaxX and PSY (Fig. 1B, S10). The amino acids in this position could affect the ability of the peptides to activate specific PSY receptor(s), as they do for XA21. Alternatively, the PSY receptor(s) may simply be able to accommodate serine or proline at this position. Further research, including the characterization of the PSY receptor(s), will help address questions of specificity and lead to a greater understanding of PSY signaling.

The study of microbial mimicry of host molecules provides insight into both host and pathogen biology, and can lead to novel strategies for disease prevention (Gardner *et al.,* 2015). Recent studies of the JA receptor have provided new insight into selective recognition of endogenous hormones. The endogenous JA receptor is sensitive to both JA-Ile and the mimic coronatine. By making a structure-guided point mutation of a single amino acid, Zhang et al. generated a modified JA receptor which has strongly reduced sensitivity to coronatine while retaining endogenous JA-Ile recognition (Zhang *et al.,* 2015). Arabidopsis with the modified JA receptor displayed enhanced resistance to coronatine producing *Pseudomonas* strains, and have a normal phenotype in the absence of infection (Zhang *et al.,* 2015). The Zhang et al. study demonstrates how understanding of bacterial mimicry of host factors can be used to engineer plants with enhanced resistance to bacterial pathogens. The findings presented in this work provide another striking example of co-evolution between the host and pathogen and provide a framework for future work directed at understanding how XA21 and the PSY receptor(s) differentially recognize RaxX and endogenous PSY peptides.

## Acknowledgments

This work was supported by NIH GM59962 and NSF IOS-1237975. The work conducted by the Joint BioEnergy Institute was supported by the Office of Science, Office of Biological and Environmental Research, of the U.S. Department of Energy under Contract No. DE-AC02-05CH11231. We thank Birgit Kemmerling (Tübingen University) for helpful discussion and for providing the *AtPSKR1/AtPSKR2/At1g72300* triple receptor mutant seeds used in this project.

## Author contributions

R.N.P., A.J., P.C.R. and W.Z. designed the research, R.N.P., A.J., W.Z. and W.F. performed experiments, J.R.D. provided resources, R.N.P, A.J. and V.S. analyzed data, R.N.P., A.J. and P.C.R. wrote the manuscript, and B.S. and W.Z. helped to revise the manuscript.

## Supplemental Information

Table S1. RaxX13 sequences from diverse Xanthomonas sources.

Table S2. Synthetic peptides used in this study.

Figure S1. Putative PSY-like proteins from Arabidopsis (At), rice (Os), banana (Ma), tomato (Sl), and wheat (Ta).

Figure S2. Comparison of the RaxX sequences from diverse bacterial strains.

Figure S3. Dose dependent activity of RaxX21-Y, RaxX21-sY, AtPSY1, and PSK on root growth of Arabidopsis *tpst-1* seedlings.

Figure S4. Sulfated RaxX21 promotes root growth in Kitaake rice.

Figure S5. Sulfated RaxX21 promotes root growth in XA21 rice.

Figure S6. Validation of the *At1g72300* mutants.

Figure S7. Addition of PSK partially blocks elf18-triggered growth inhibition in Arabidopsis seedlings, whereas RaxX21-sY and AtPSY1 do not.

Figure S8. PXO99 strain lacking RaxX is impaired in virulence.

Figure S9. Sequence similarity of RaxX from *Xoo* and *Xoc* with selected rice PSYs.

Figure S10. Comparison of RaxX and PSY peptides from various species.

